# The research progress of p53 tumour suppressor activity controlled by Numb in triple-negative breast cancer

**DOI:** 10.1101/463802

**Authors:** Jie Xian, Yu Cheng, Xue Qin, Yequn Luo, Yijia Cao, Youde Cao

## Abstract

Numb is known as a cell fate determinant as it determines the direction of cell differentiation by asymmetrically partitioning at mitosis. It has been shown to be a tumor suppressor, and there is frequent loss of Numb expression in breast cancer. Numb enters in a tricomplex with p53 and the E3 ubiquitin ligase HDM2 (also known as MDM2), thereby preventing ubiquitination and degradation of p53. The aim of this study was to investigate the expression and migration of Numb, HDM2 and p53 proteins in the membrane, cytoplasmic and nuclear fractions of MCF-10A cells and MDA-MB-231 cells. We extracted the cell fractions to detect changes of these three protein levels after re-expression of NUMB in the basal-like triple-negative cell line MDA-MB-231 and knocking down NUMB in the normal mammary epithelial cell line MCF-10A. Our results show that Numb protein can migrate from cytoplasm to nucleus of the MDA-MB-231 cells. Numb protein regulates p53 levels in nucleus of MCF-10A cells and MDA-MB-231 cells, and the Numb levels is positively correlated with p53 levels. We have a finding on HDM2 protein that after knocking down NUMB in MCF-10A cells, it was remarkably reduced in the membrane fraction of NUMB knockdown cells, but its mRNA levels was significantly increased. We also examined the Numb expression in 125 patients with triple-negative breast cancer, 61(48.8%) displayed deficient or reduced expression of Numb. The percent of Ki67>14% in retained Numb group was significantly lower than that in the reduced and deficient Numb group (86.00% vs. 98.40%, P=0.0171).

## Introduction

Numb protein was first discovered in the Drosophila sensory neural precursor cells (SOP). It is found that the asymmetrically partitioning of Numb protein in the division of SOP determines the differentiation direction of daughter cells, thereby it is called the cell fate determiner (Rhyu, 1994; Berdnik, 2002; Cayouette, M. & Raff, M, 2002). Breast cancer is a common malignant tumor in women, and its incidence rate is increasing every year. Triple-negative breast cancer (TNBC) is a type of malignant breast cancer, which represents a subgroup of breast cancers that is negative for estrogen receptors (ER), progesterone receptors (PR), and human epidermal growth factor receptor 2 (HER2) (Foulkes WD, 2010). In breast cancers there is frequent loss of Numb expression. Immunohistochemical staining of pathological sections of 241 breast cancer patients revealed that deficient or reduced Numb expression up to 44% in the 25 cases of the basal-like triple-negative breast cancer groups, and its loss or reduction was correlated to the degree of the tumor malignancy, patient prognosis and five-year survival rate (Pece, S. et al, 2004). We performed immunohistochemical staining on paraffin sections of 125 patients with triple-negative breast cancer, diagnosed in the Clinical pathology diagnosis center of Medical University of Chongqing between 2012 and 2017 to detect the expression of Numb The results show that 61(48.8%) displayed deficient or reduced expression of Numb, it further prove that there was a significant positive correlation between Numb expression (deficient and reduced vs. retained) and ER and PR status (Pece, S. et al, 2004). The percent of Ki67>14% in retained Numb group was significantly lower than that in the reduced and deficient Numb group (P=0.0171), but there is no significant difference in age, tumor size, lymph node status, and histological type (between the retained Numb group and (reduced and deficient) Numb group.

The mammary epithelium is composed of the inner layer of luminal epithelial cells and the outer layer of myoepithelial cells with mesenchymal characteristics. Numb and NumbL (Numb-like) are abundantly expressed in mammary muscle epithelial cells and peak in mice during pregnancy. It was found that knocking out mice Numb and NumL will damage mammary muscle epithelial cells and promote its epithelial-to-mesenchymal transition (ETM) leading to lactation failure (Yue Zhang, 2016). The results of this study further demonstrated that the loss of Numb expression is correlated to the occurrence of breast cancer. The reduction of Numb expression also affects cell cycle proteins, thus accelerating the transformation of G1/S phase and promoting the proliferation of tumor cells (Lucia Di Marcotullio, 2006). Mutation and reduction of the tumor suppressor p53 in breast cancer is common.HDM2 is an oncoprotein that can degrade P53, resulting in shorter half-life and reduced activity of p53 (Kubbutat, 1997; Vassilev, 2004; Momand, 1992; Haupt, 1997). Numb protein interacts with HDM2, preventing its ubiquitination and degradation of p53, thus maintaining the activity and stability of p53 (Ivan N, 2008). However, due to the different distribution of Numb, HDM2 and p53 in the cell, the specific mechanism of how they interact is not clear.

The aim of the present study was to further study the migration and the expression of Numb, HDM2 and p53 in the cell membrane, cytoplasm and nucleus of MCF-10A cells and MDA-MB-231 cells. The results showed that Numb and HDM2 proteins distribute in the cell membrane, cytoplasm and nucleus, while p53 was mainly distributed in the nucleus of MCF-10A cells and MDA-MB-231 cells. The expression of these three proteins was higher in MCF-10A cells than MDA-MB-231 cells. Next, we re-expressed NUMB in MDA-MB-231 cells and knocked down NUMB in MCF-10A cells by using short interfering RNA (siRNA), respectively, and then detect changes of the three proteins in the membrane, cytoplasmic and nuclear fractions of cells. The results indicated that either re-expressed NUMB in MDA-MB-231 cells or knocked down NUMB in MCF-10A cells, the protein level of Numb was remarkably changed in the nuclear fraction, meanwhile no significant change of HDM2 was observed. NUMB– EGFP-transfected MDA-MB-231 cells displayed an approximately twofold-higher level of p53 and Numb in the nucleus, and HDM2 levels was no significant change. These results prove that in nucleus of the MDA-MB-231 cells, Numb is a key factor regulating the p53 levels, which can inhibit the degradation of p53 by HDM2. However, there was unchanged of Numb in the cell membrane and cytoplasm. These results prove that Numb protein can migrate from the cytoplasm to nucleus of NUMB–EGFP-transfected MDA-MB-231 cells.

After knocking down NUMB in MCF-10A cells, the protein levels of HDM2 evidently decreased, but HDM2 mRNA levels significantly increased. Further analysis showed that HDM2 in the cell membrane was significantly reduced, and there was no change in both the cytoplasm and the nucleus of NUMB knockdown cells. In the unclear fraction of NUMB knockdown cells, the expression of Numb and p53 was markedly increased, and HDM2 protein levels was unchanged. The p53 mRNA levels was not changed. Above all, these results prove that Numb regulate p53 levels in the nucleus of MCF-10A cells and MDA-MB-231 cells, and it was positively correlation with p53 levels.

## Results

### Numb expression in Normal mamary tissue and Triple-negative breast cancers

We performed immunohistochemical staining on paraffin sections of 125 patients with triple-negative breast cancer to detect the expression of Numb. Of the 125 patients evaluated for Numb expression, 64 displayed retained expression (score 2; 51.2%), 43 displayed reduced expression (score 1; 34.4%) and 18 were Numb deficient (score 0; 14.4%). The associations between Numb expression and patient and tumor characteristics in the studied cohort are summarized in Table 1. The percent of Ki67>14% in retained Numb group was significantly lower than that in the reduced and deficient Numb group (86.00% vs. 98.40%, P=0.0171) (Table 1). There is no significant difference in age (χ2 test; P=0.2796), tumor size (χ2 test; P=0.5911), lymph node status (χ2 test; P=0.6091), and histological type (Fisher’s exact test; P=0.5762) between the retained Numb group and (reduced and deficient) Numb group.

**Table 1.**
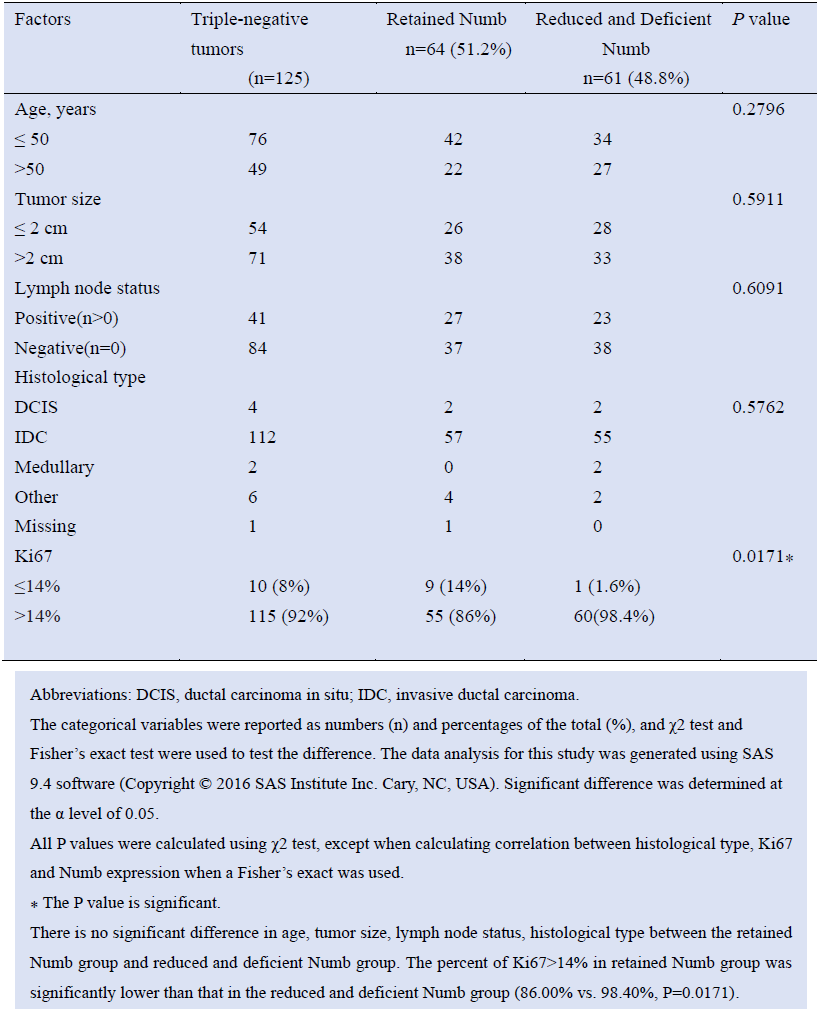
Associations between Numb expression and patient and tumor characteristics.

### The localization and expression of Numb, HDM2 and p53 in MCF-10A and MDA-MB-231 cell lines

Cellular immunofluorescence experiment was used to locate the two proteins of Numb and HDM2 in MCF-10A and MDA-MB-231 cells. The results showed that Numb was mainly distributed in the membrane fraction of both cell lines, and weakly fluorescence was detected in the nucleus (Fig 2A). HDM2 is distributed in the cell membrane, cytoplasm and nucleus of the MCF-10A cells and MDA-MB-231 cells. (Fig 2A). Western-blot experiments indicated that the protein levels of Numb, p53 and HDM2 in MCF-10A cells were higher than those in MDA-MB-231 cells, and this was the same trend as q-PCR results (Fig 2B and F). We further isolated and extracted the cell fractions. Western-blot experiments showed that Numb and HDM2 proteins were distributed in the cell membrane, cytoplasm and nucleus, while p53 was mainly distributed in nucleus of MCF-10A cells (Fig 2B). Numb, HDM2 and p53 proteins in the nuclear fraction were all higher in MCF-10A cells than MDA-MB-231 cells (Fig 2B). In the membrane fraction, the protein level of Numb and HDM2 were higher in MCF-10A cells than MDA-MB-231 cells, and p53 protein was not detected (Fig 2B). In the cytoplasmic fraction, the expression of Numb in MCF-10A cells was evidently higher than in MDA-MB-231 cells, but there was no difference of HDM2 and p53 protein levels (Fig 2B).

**Figure. 1.**
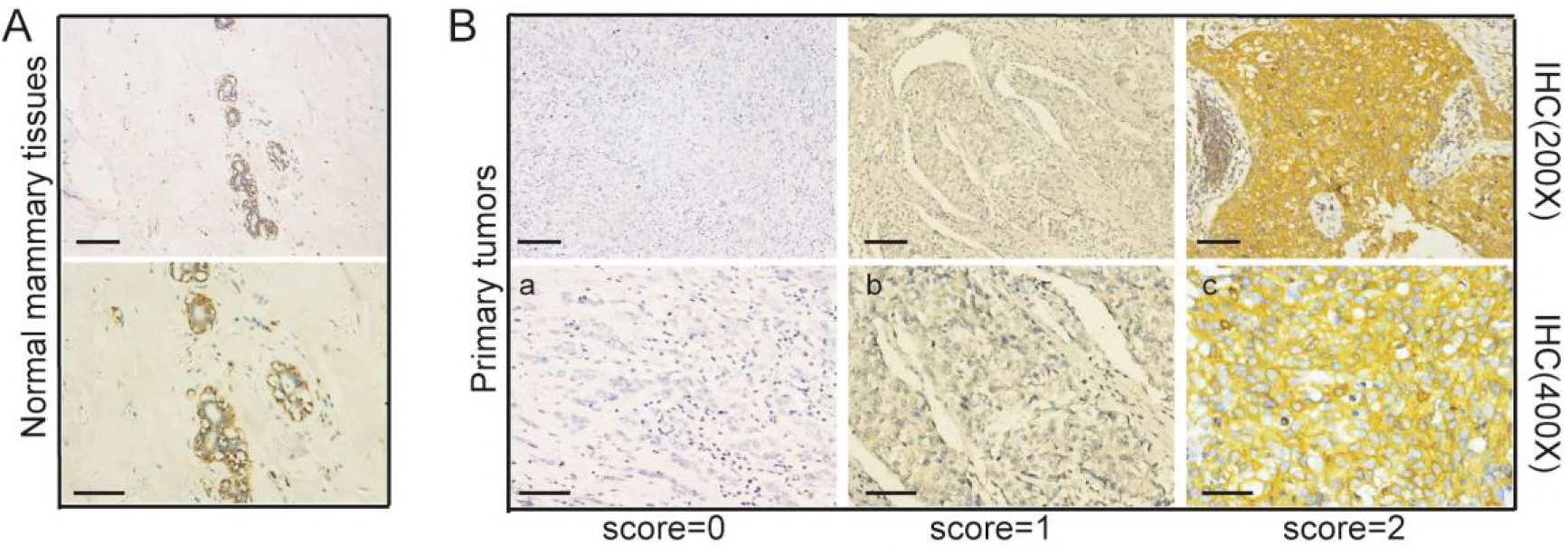
Numb expression in Normal breast tissue and Triple-negative breast cancers. A The expression of Numb in Normal mammary tissue detected by immunohistochemistry. B Examples of a retained (score 2, <10% positive tumor cells), b reduced (score 1, 10–50%positive tumor cells) and c deficient (score 0, >50% positive tumor cells), expression of Numb.

**Figure2.**
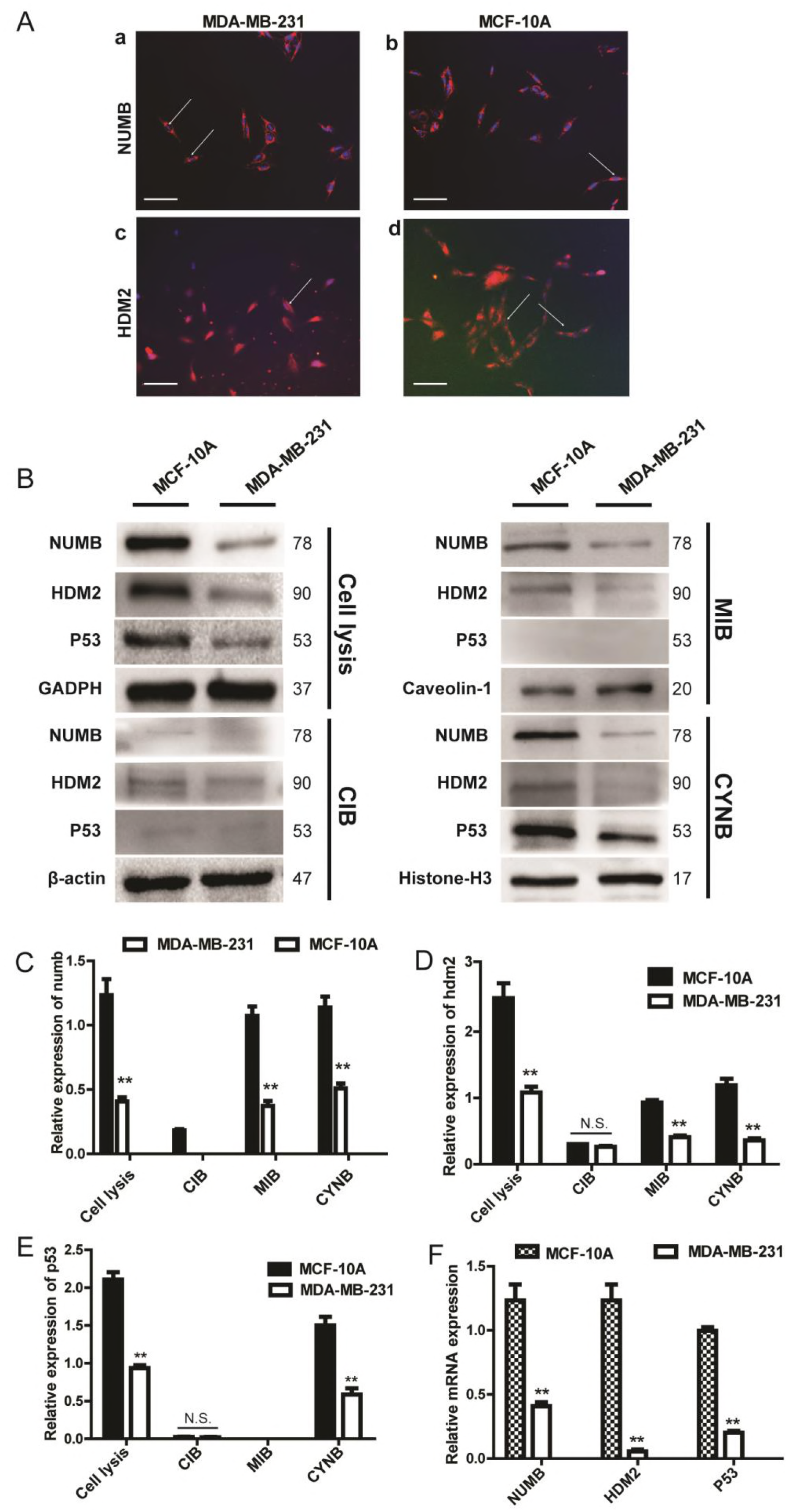
The localization and expression of Numb, HDM2 and p53 in MCF-10A and MDA-MB-231. Cell lysis, total protein; MIB, membrane fraction; CIB, cytoplasmic fraction; CYNB, nuclear fraction A Immunofluorescence staining of NUMB and HDM2.(a)The localization of NUMB in MDA-MB-231 (X400);(b)The localization of NUMB in MCF-10A (X400);(c)The localization of HDM2 in MDA-MB-231 (X400);(d)The localization of HDM2 in MCF-10A(X200) Scale bar represented 50μm B The protein expression of NUMB,HDM2 and p53 in different cell fractions of MDA-MB-231 and MCF-10A was determined by Western blot. C, D, E Quantitative analysis of NUMB, HDM2 and p53 expression in different cell fractions of MDA-MB-231 and MCF-10A ( Mean ± SD, n = 3, **P < 0.01). F Quantitative RT–PCR detection of *NUMB, HDM2* and *p53* levels in MDA-MB-231 and MCF-10A.

Above all, the Numb levels were remarkably higher in all cell fractions of MCF-10A cells than MDA-MB-231 cells (Fig 2C). The HDM2 levels of the cytoplasmic fraction was no different between these two cell lines, but it was higher expression in the cell membrane and nucleus of MCF-10A cells than MDA-MB-231 cells (Fig 2D). The expression of p53 in the nuclear fraction was higher remarkably in MCF-10A cells than MDA-MB-231 cells. In the cytoplasmic fraction, p53 expression was very low and there was no significant difference between MDA-MB-231 cells and MCF-10A cells. In addition, p53 was not expressed in the cell membrane (Fig 2E).

### The Numb, HDM2 and p53 levels in different cell fractions of NUMB-EGFP-transfected MDA-MB-231 cells

The above experimental data suggested that Numb was highly expressed in MCF-10A, while relatively low expressed in the basal-like cell line MDA-MB-231. Therefore, we re-expression *NUMB* in MDA-MB-231 cells by transfecting with NUMB-EGFP plasmid. Further, after separating different cell fractions for Western-blot experiments, it was found that the Numb and p53 proteins was significantly increased in the nuclear fraction of NUMB-EGFP-transfected MDA-MB-231 cells, while HDM2 was not evidently changed (Fig 3A). However, in the fraction of membrane and cytoplasm, Numb, HDM2 and p53 have not been affected all (Fig 3A). Meanwhile, there was no markedly change of *HDM2* and *P53* in the mRNA level after re-expression of *NUMB* in MDA-MB-231 (Fig 3E).

**Figure 3.**
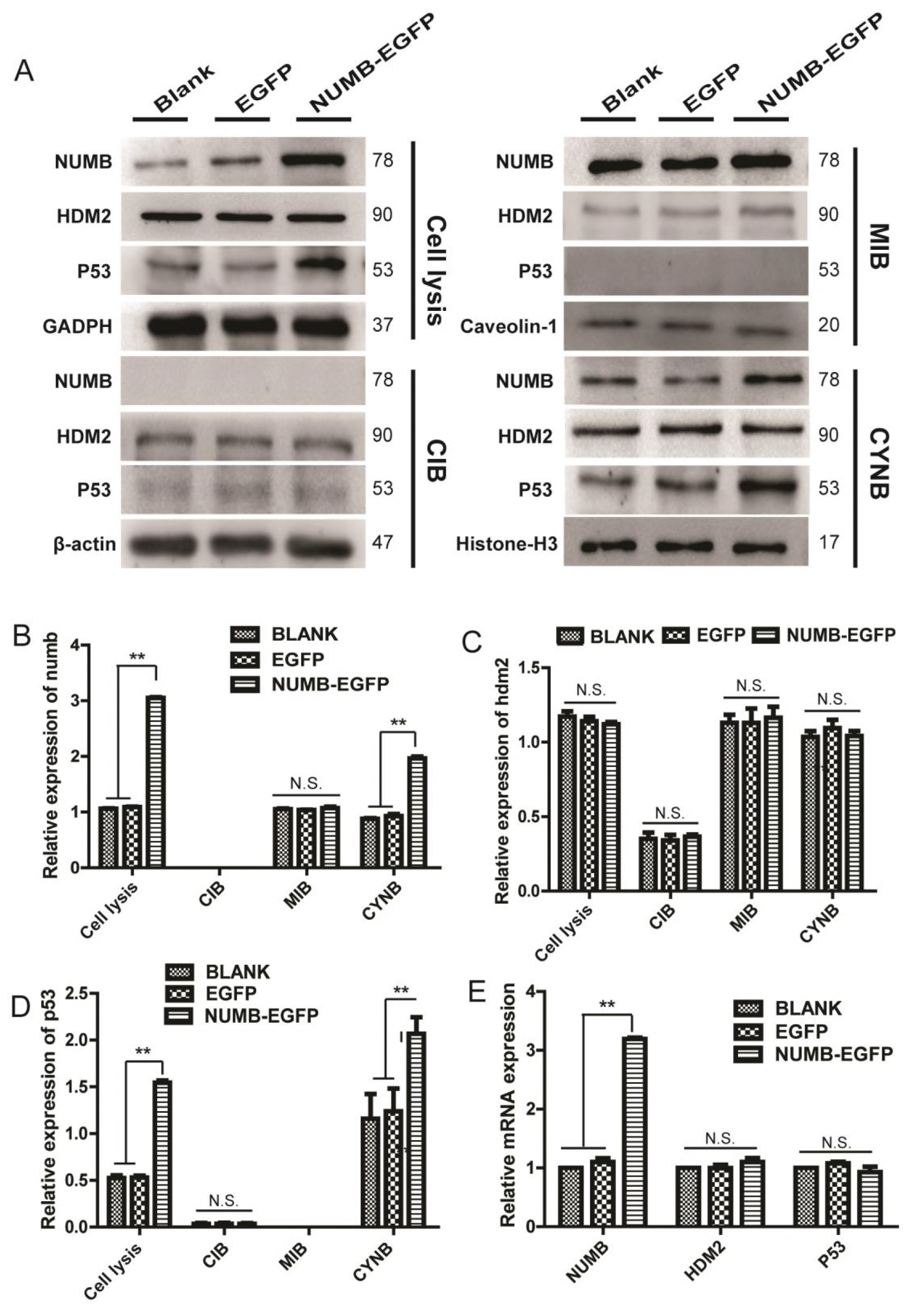
Effects of *NUMB* re-expression on Numb, HDM2 and p53 proteins in different cell fractions of MDA-MB-231 cells. A The protein expression of Numb, HDM2 and p53 in the membrane, cytoplasmic and nuclear fractions of NUMB-EGFP-transfected MDA-MB-231 cells was determined by Western blot. B, C, D Quantitative analysis of Numb, HDM2 and p53 proteins in different cell fractions of NUMB-EGFP-transfected cells. (Mean±SD, n=3, **P<0.01, vs blank group or EGFP group). E Quantitative RT–PCR detection of *NUMB*, *HDM2* and *p53* levels in NUMB-EGFP-transfected MDA-MB-231 cells.

Above all, the expression of Numb in the membrane fraction was unchanged significantly, in the cytoplasmic fraction was still no expression, in the nuclear fraction was increased remarkably after re-expression *NUMB* in MDA-MB-231 cells (Fig 3B). The HDM2 levels was no changed in any fractions of NUMB-EGFP-transfected MDA-MB-231 cells (Fig 3C). The p53 levels was significantly increased in the nuclear fraction, and no expression in the membrane fraction and the cytoplasmic fraction of the NUMB-EGFP-transfected MDA-MB-231 cells (Fig 3D).

### Effects of *NUMB* knockdown on Numb, HDM2 and p53 expression in different cell fractions of MCF-10A cells

Transfection with NUMB siRNA decreased the Numb protein expression in MCF-10A cell line, the protein level of HDM2 and p53 decreased accordingly (Fig 4A). The q-PCR indicated that the mRNA level of *HDM2* was significantly increased, while the *p53* was not evidently changed (Fig 4E). Further, after separating the cell fractions for Western-blot experiments, it was found that the protein level of Numb and p53 was both decreased in the nuclear fraction, while HDM2 was unchanged (Fig 4A). In the membrane fraction of MCF-10A, Numb did not change significantly, while the HDM2 protein decreased remarkably, and p53 was not detected (Fig 4A). The Numb, HDM2, and p53 levels were no significantly difference in the cytoplasmic fraction. (Fig 4A).

**Figure 4.**
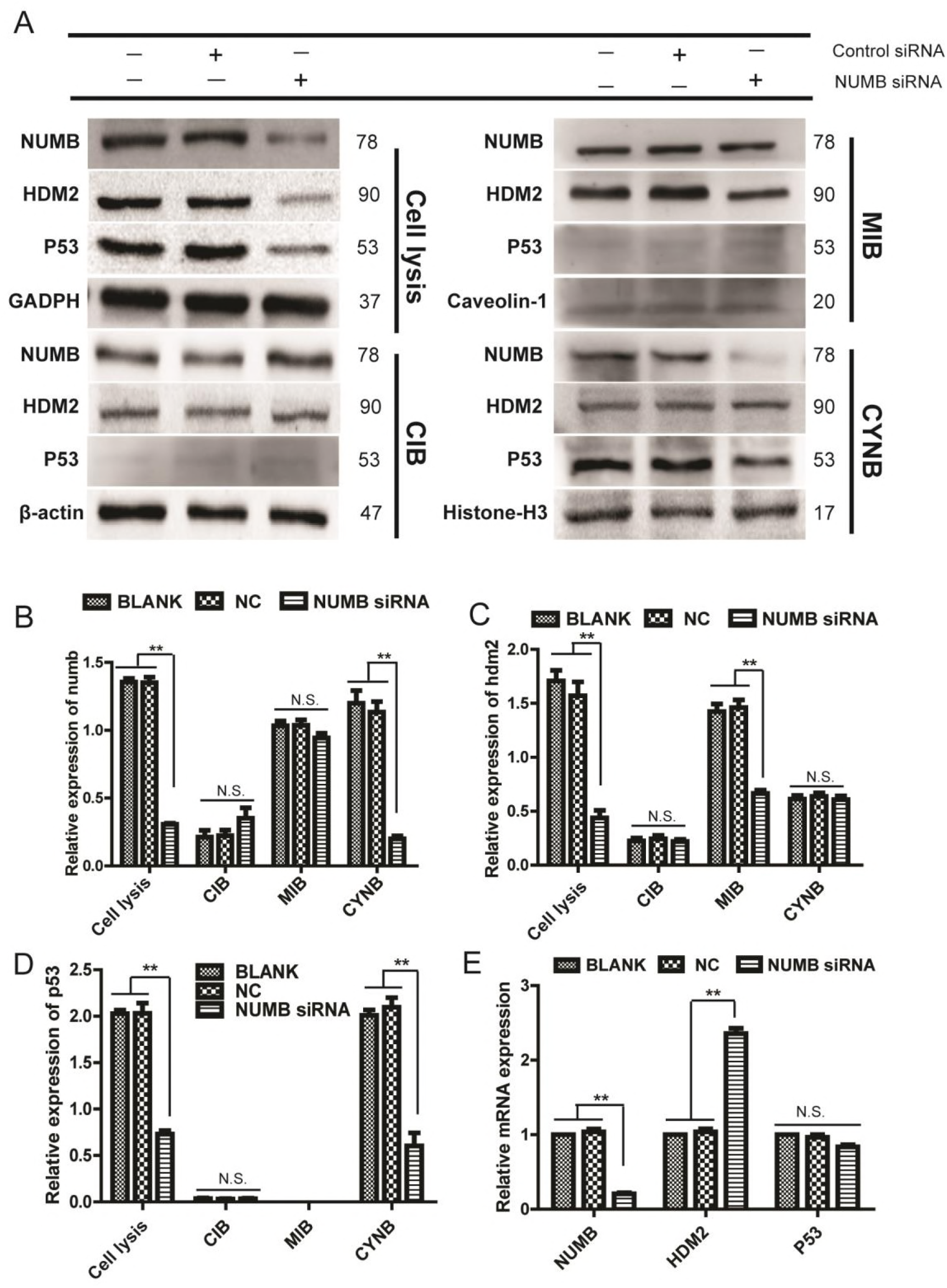
Effects of *NUMB* knockdown on Numb, HDM2 and p53 proteins in different cell fractions of MCF-10A cells. A The protein expression of Numb, HDM2 and p53 in the membrane, cytoplasmic and nuclear fractions of Numb knockdown MCF-10A cells was determined by Western blot. B,C,D Quantitative analysis of Numb, HDM2 and P53 proteins in different cell fractions of Numb knockdown cells (Mean±SD, n=3 **P<0.01, vs blank group or NC group). E The mRNA expression of *NUMB,HDM2* and *p53* in Numb knockdown cells was determined by q-PCR.

Above all, the Numb levels was decreased evidently in the nuclear fraction after tampering with *NUMB* in MCF-10A cell line, and it unchanged significantly in the membrane fraction and the cytoplasmic fraction (Fig 4B). The expression of HDM2 was decreased significantly in the membrane fraction of MCF-10A, and it was unchanged in the cytoplasmic and nuclear fraction (Fig 4C). The p53 levels was evidently increased in the nucleus of MCF-10A cells, and it was unchanged in the cytoplasm. It was no expression in the cell membrane (Fig 4D).

### Discussion

A higher percentage of the tumors from the triple-negative (SR-/HER2-) subgroup displayed reduced or deficient Numb expression compared to tumors in the other subgroups (44% [11/25] of the basal-like) (Karin, 2010). So we performed immunohistochemical staining on paraffin sections of 125 patients with triple-negative breast cancer to detect the expression of Numb. The results showed that 64 (51.2%) cases displayed retained expression, 61(48.8%) displayed reduced or deficient expression. This illustrates that reduced or deficient Numb expression in triple-negative breast cancer is common, nearly 50%. Accordingly, we focus on the relationship between Numb and triple-negative breast cancer, and further investigate the molecular mechanism of Numb to promote the development of disease. It will discover potential intervention targets for the disease. We further analyzed the associations between Numb expression and patient and tumor characteristics. The percent of Ki67>14% in retained Numb group was significantly lower than that in the reduced or deficient Numb group (86.00% vs. 98.40%, P=0.0171). This finding is also in agreement with previously published data on the inverse correlation between Numb expression levels and indicators of aggressive disease (Pece S, 2004; Karin, 2010).

It has been reported that Numb inhibits the degradation of p53 by binding to HDM2 in MCF-10A cell line, thus maintaining the activity and stability of p53 (Ivan N, 2008). Studies have shown that Numb protein is mainly distributed in cell membrane and cytoplasmic fraction, while HDM2 and p53 are by-and-large nuclear proteins (). But where the three proteins interact and their specific mechanisms of action are still unknown. WB and cell immunofluorescence staining results show that Numb and HDM2 proteins were distributed in the membrane, cytoplasmic and nuclear fractions of MCF-10A, while p53 was mainly distributed in nucleus. Numb is not distributed in cytoplasm of MDA-MB-231, HDM2 is distributed in all cell fractions, and p53 is mainly distributed in nucleus. We detected the, expression and transcription levels of *NUMB, HDM2* and *p53* in McF-10A and MDA-MB-231, and found the higher levels of these three proteins in MCF-10A than in MDA-MB-231. In the cell membrane, the protein levels of Numb and HDM2 was significantly higher in MCF-10A than in MDA-MB-231 cells, while p53 was not detected. In the cytoplasmic fraction, the expression of Numb in MCF-10A was higher than in MDA-MB-231, while there was no difference of HDM2 and p53 levels. In the nuclear fraction, the expression of Numb, HDM2 and p53 was all higher in MCF-10A than in MDA-MB-231 cells. HDM2 is an oncogene and can degrade p53, but we found that in the normal breast epithelial cell line MCF-10A, the HDM2 levels was remarkably higher than in the basal-like triple-negative breast cancer cell line MDA-MB-231. Surprisingly, the protein levels of p53 in MCF-10A cells was also evidently higher than in MDA-MB-231 cells. This may be due to Numb was also highly expressed in MCF-10A cells. Moreover, In *NUMB* knockdown MCF-10A cells, the level of HDM2 protein on the cell membrane was significantly reduced after the numb protein in the nucleus was significantly reduced. It is suggested that tumor suppressor Numb regulates oncoprotein HDM2.The Numb expression was strongly decreased in the basal-like triple-negative breast cancer cell line MDA-MB-231, so we re-expressed *NUMB* in MDA-MB-231 cells with the NUMB-EGFP plasmid and then detected the changes of Numb, HDM2 and p53 levels in each cell fraction. The results showed that after re-expression of *NUMB*, there was no significant change of the three proteins in the cytoplasmic and membrane fraction of MDA-MB-231 cells. However, in the nuclear fraction, Numb and p53 protein were significantly increased, while HDM2 levels was not markedly changed. q-PCR showed that the transcriptional levels of *HDM2* and *p53* was not changed after re-expression of *NUMB*. Therefore, we assumed that the increasing of p53 protein is caused by the increasing of Numb protein in the nucleus, and the expression of Numb in the nucleus was positively correlated with p53 levels. This finding is also in agreement with previously published data that NUMB-mediated regulation of p53 at the post-transcriptional level in MCF-10A cells (Ivan N, 2008).

The above results showed that re-expression *NUMB* in MDA-MB-231 cells, Numb levels has not changed in the membrane and cytoplasmic fractions, it only increased in the nucleus. It indicated that Numb can migrate into the nucleus from the cytoplasm of MDA-MB-231 cells to regulate the of p53 levels by binding with HDM2.

It has been reported that after knocking down *NUMB* in MCF-10A cells, the protein levels of Numb, HDM2 and p53 were all decreased remarkably, but *p53* mRNA levels did not change. This indicates that Numb regulates p53 levels at post-transcriptional levels (Ivan N, 2008). We also knocked down *NUMB* in MCF-10A cells by using siRNA, and then isolated cell fractions for analysis. We found that there was no significant reduction of Numb in the membrane and cytoplasmic fractions, but a significant reduction in the nuclear fraction of *NUMB* knockdown cells. In the nuclear fraction, the expression of NUMB was remarkably increased, HDM2 levels was not evidently changed, and p53 levels was markedly reduced. It was further demonstrated that whether in MDA-MB-231 cells or in MCF-10A cells, Numb regulate the p53 levels in the nucleus where the protein levels of Numb is positively correlated with p53 protein levels. This finding is also in agreement with previously published data that the reduction in p53 levels was caused by loss of Numb, by means of HDM2 (Ivan N, 2008).

Whether re-expression of *NUMB* in MDA-MB-231 or interference with *NUMB* in MCF-10A, there was no evidently change of Numb in the cell membrane and the cytoplasm, but there was a markedly change in the nucleus, which further indicates that the Numb protein exists in the nucleus and plays an important role in regulating the p53 levels.

We also have a new finding on HDM2 protein that after knocking down *NUMB* in MCF-10A cells, it was significantly reduced in the membrane fraction, but its mRNA levels was significantly increased. Therefore, we think the reduction of HDM2 is caused by the regulation of post-transcriptional levels, and then HDM2 protein levels reduction lead to the increase of mRNA levels. But we have no idea the mechanism of how HDM2 protein in the cell membrane remarkably induced after Numb decreased in nucleus of MCF-10A cells, and whether the decrease of HDM2 in the membrane fraction causes the affection in the downstream pathway. We speculate that a significant increase of HDM2 in the transcription level may be due to feedback regulation caused by the HDM2 decrease in cell membrane or due to the significant reduction of Numb and p53 levels in the nucleus of *NUMB* knockdown MCF-10A cells.

## Materials and Methods

### Clinical samples

We collected archival formalin-fixed paraffin-embedded (FFPE) mammary tissue specimen of 100 Cases with Triple-negative Breast Cancer, diagnosed in the Clinical pathology diagnosis center of Medical University of Chongqing between 2012 and 2017. The Numb status was attributed to the tumors by measuring the levels of Numb expression by IHC. Normal mammary tissues displayed strong and homogeneous Numb staining in the luminal layers (IHC score 2) (see also Pece et al, 2004; Karin et al, 2010). Tumors were classified on an IHC scale from 0 to 2 (score 2, <10% positive tumor cells; score 1, 10–50%positive tumor cells; score 0, >50% positive tumor cells expression of Numb).

### Cell culture, plasmids and reagents

MCF10A and MDA-MB-231 cell lines were obtained from the American Type Culture Collection. MCF10A cells were cultured in MEBM (CC-3151, Lonza, Basel, Switzerland) supplemented with MEGM®SingleQutos® (CC-4136, Lonza, Basel, Switzerland), 100 ng/mL cholera toxin. MDA-MB-231 cells were cultured in DMEM containing 10% FBS(Corning). The plasmid of plRES2-EGFP-NUMB (PPL00760-2a) was purchased in Public Protein/Plasmid Library. We used Lipofectamine2000 (180423, GenePharma, shanghai, China) to transfect MDA-MB-231 cells.

### Immunohistochemistry

Serial sections were cut from paraffin blocks, deparaffinized with xylene and then rehydrated in a graded ethanol series. For antigen retrieval, the samples were microwaved for 14 min in a citrate buffer (pH = 6). Subsequently, the sections were treated with 3% hydrogen peroxide for 15 min. All sections were incubated with Anti-NUMB monoclonal antibody(1:100, ab14140; Abcam, Cambridge, MA, USA) overnight at 4°C. These antibodies were detected using a biotinylated secondary antibody (PV-9001, Beijing Sequoia Jinqiao, Beijing, China) labeled with streptavidin-horseradish peroxidase (HRP) and a DAB staining kit (KIT-5020, Maixin Biotechnology, Fuzhou, China). As a negative control, the primary antibody was substituted with PBS. The same tonsil tissues were used as positive controls for both antibodies.

### Immunofluorescence

Cells were seeded onto glass coverslips in 24-well plates, washed with PBS, fixed in 4% paraformaldehyde for 20 min at room temperature, permeabilized with 0.1% Triton X-100. Cells incubated in blocking buffer (PBS with 5% BSA) for 30 min and then with primary antibody overnight in PBS at 4°C. And then, we incubated samples with secondary antibodies (1:100, bs-0296G-Cy3/bs-0295G-FITC; Bioss, Beijing, China) for 1h and stained with DAPI for 10 min at room temperature.

### Cell Fractionation expriments and immunoblotting

We obtained the cytoplasmic fraction, membrane and organelle fraction, cytoskeletal and nuclear fraction of cells by lysing in Cell Fractionation Kit (#9038, Cell Signaling Technology, Danvers, MA, USA). Whole cell proteins were extracted by lysing in RIPA buffer. The protein lysate was separated using 10% sodium dodecyl sulfate-polyacrylamide gel electrophoresis (SDS-PAGE) and electrotransferred onto a polyvinylidene fluoride (PVDF) membrane. After blocking in 5% bovine serum albumin (BSA), the membranes were incubated with primary antibodies overnight at 4°C. The following antibodies were used: mouse monoclonal anti–NUMB (1:1000,ab14140; Abcam, Cambridge, MA, USA), mouse monoclonal anti-MDM2[2A10] (1:50,ab16895; Abcam, Cambridge, MA, USA), Rabbit Polyclonal anti-P53(1:1000, #9282; Cell Signaling Technology, Danvers, MA, USA), Rabbit Monoclonal anti-Histone H3(D1H2) (1:2000, #4499; Cell Signaling Technology, Danvers, MA, USA), Rabbit Polyclonal anti-Caveolin-1(1:1000, bs-1453R; Bioss, Beijing, China) and mouse anti-GAPDH (1:1000, 60004 –1-Ig; Proteintech Group, Wuhan, China) antibodies. The membranes were washed and incubated with anti-rabbit or anti-mouse secondary antibody (1:5000, SSA004/SSA007; Sino Biological Inc., Beijing, China) for 2h at room temperature. Finally, antigen–antibody complexes were detected using an electrochemiluminescence (ECL) Western blotting detection reagent.

### RNA isolation, q-PCR and siRNA

Total RNA was isolated with RNAiso Plus (TaKaRa, Kusatsu, Shiga, Japan) and reverse transcribed with PrimeScript® RT reagent Kit with gDNA Eraser (Perfect Real Time) (TaKaRa, Kusatsu, Shiga, Japan). q-PCR quantification was performed using SYBR® Premix Ex Taq™ II (Tli RNaseH Plus) (TaKaRa, Kusatsu, Shiga, Japan) on the CFX96 Real-Time PCR Detection System (Bio-Rad Laboratories Inc, Hercules, California, USA). The following primers were used for aquantitative q-PCR: NUMB: sense primer, 5′-GGACACAGGTGAAAGGTTGAGC-3′, and anti-senseprimer,5′-AGTGGCTGTTGTGACACGGAAT-3′; MDM2: sense primer, 5′-CTACAGGGACGCCATCGAATC-3′, and anti-sense primer, 5′-TGAAGTGCATTTCCAATAGTCAGC-3′; P53: sense primer, 5′-TGCGTGTTTGTGCCTGTCCT-3′, and anti-sense primer, 5′-AGTGCTCGCTTAGTGCTCCCT-3′; GAPDH: sense primer, 5′-CTTTGGTATCGTGGAAGGACTC-3′, and anti-sense primer, 5′-GT AGAGGCAGGGATGATGTTCT-3′. The reaction conditions were 95°C for 30 s, followed by 40 cycles at 95°C for 5 s and 58°C for 30 s. The housekeeping gene GAPDH was used for normalization, and all reactions were performed in triplicate. The relative mRNA expression was analyzed using the 2-δΔCt method. For siRNA experiments, delivery of siRNA oligos was achieved using siRNA-mate (180426, GenePharma, shanghai, China). The targeted sequences were as follows: Numb siRNA, GGUUAAGUACCUUGGCCAUTT; AUGGCCAAGGUACUUAACCTT. (5′ to 3′)

### Statistical analysis

The Student’s t test (two-tailed) was used to determine statistically the significance of differences between groups. P< 0.05 was considered statistically significant. NUMB expression intensities in human breast cancer samples were analyzed by χ2 test and Fisher’s exact test. The data analysis for this study was generated using SAS 9.4 software (Copyright © 2016 SAS Institute Inc. Cary, NC, USA). Significant difference was determined at the α level of 0.05.

## Acknowledgements

We thank all the breast cancer patients who donated their samples for research. We also thank Manran Liu for MCF-10A cells; the Chenglong Wang for histopathological analysis. This work was supported by grants from the Chongqing science and technology commission to YDC and JX.

## Author contributions

JX, XQ performed experimental work and analyzed data. YC, YL, JX performed data analysis, and wrote the manuscript. YC planned and supervised the project, provided samples and supervised the histopathological analysis.

